# Genotype-covariate correlation and interaction disentangled by a whole-genome multivariate reaction norm model

**DOI:** 10.1101/377796

**Authors:** Guiyan Ni, Julius van der Werf, Xuan Zhou, Elina Hyppönen, Naomi R. Wray, Hong Lee

## Abstract

The genomics era has brought useful tools to dissect the genetic architecture of complex traits. We propose a reaction norm model (RNM) to tackle genotype-environment correlation and interaction problems in the context of genome-wide association analyses of complex traits. In our approach, an environmental risk factor affecting the trait of interest can be modeled as dependent on a continuous covariate that is itself regulated by genetic as well as environmental factors. Our multivariate RNM approach allows the joint modelling of the relation between the genotype (G) and the covariate (C), so that both their correlation (association) and interaction (effect modification) can be estimated. Hence we jointly estimate genotype-covariate correlation and interaction (GCCI). We demonstrate using simulation that the proposed multivariate RNM performs better than the current state-of-the-art methods that ignore G-C correlation. We apply the method to data from the UK Biobank (N= 66,281) in analysis of body mass index using smoking quantity as a covariate. We find a highly significant G-C correlation, but a negligible G-C interaction. In contrast, when a conventional G-C interaction analysis is applied (i.e., G-C correlation is not included in the model), highly significant G-C interaction estimates are found. It is also notable that we find a significant heterogeneity in the estimated residual variances across different covariate levels probably due to residual-covariate interaction. Using simulation we also show that the residual variances estimated by genomic restricted maximum likelihood (GREML) or linkage disequilibrium score regression (LDSC) can be inflated in the presence of interactions, implying that the currently reported SNP-heritability estimates from these methods should be interpreted with caution. We conclude that it is essential to correctly account for both interaction and correlation in complex trait analyses and that the failure to do so may lead to substantial biases in inferences relating to genetic architecture of complex traits, including estimated SNP-heritability.

## INTRODUCTION

Variation in complex traits between people is determined both by genetic and non-genetic factors. The non-genetic component will include known environmental risk factors, but also unknown factors that are characterised by stochastic variation. The interplay between genetic and identifiable environmental factors has long been a topic of research interest^1^, since the identification of genotype-environment interactions has the potential to inform on health interventions to overcome genetic predisposition to disease. However, many so-called environmental risk factors (e.g., smoking, alcohol consumption, stressful life events, educational attainment) are themselves complex traits whose variation also reflects both genetic and non-genetic factors. For example, the relationship between smoking and body mass index (BMI) is complex, i.e. common causal genetic variants have biological effects on both traits (pleiotropy or genetic correlation)^2^ while BMI is also affected by smoking status (interaction)^3; 4^. The relationship between smoking and BMI is a good example for a complex association which can be best modelled using a framework that can account both for genotype-covariate correlation and interaction (GCCI).

Both correlation (‘association’) and interaction (‘effect modification’) are fundamental in biology^5-7^, but it is critical to distinguish between them because their biological mechanisms differ, as do their implications. This association/interaction problem has been well posed in the classical twin study approach^8^, showing that association and interaction can be disentangled and correctly estimated with an appropriate model and sufficient data. Unfortunately, large well-powered data sets with measures on multiple family members are limited. However, genome-wide association studies (GWAS) now provide different type of genetically informative data to investigate GCCI. The genomic era has brought useful tools to dissect the genetic architecture of complex traits, where genetic variance and covariance can be estimated based on genome-wide single nucleotide polymorphisms (SNPs) genotyped in large-scale population samples. The increased availability of sufficiently powered data sets, with information on measured genetic and non-genetic risk factors, motivates the need to develop appropriate statistical tools for GCCI analysis.

The reaction norm model (RNM) has been developed and applied to GCCI analyses in ecology^9^ and agriculture^10; 11^. The RNM allows environmental exposures to be modelled such that the genetic effects of a trait can be fitted as a nonlinear function of a continuous environmental gradient. The possible modulators of the phenotypes of the trait are not limited to environmental exposures, but can include any covariates, regulated by environmental and genetic factors, which are shared with the phenotypes. In other words, the genetic effect, and therefore the phenotype, of one trait often depends on the phenotype of another trait. This can be modelled by introducing dependence between the phenotype and the covariate, where the covariate represents the phenotype of the modulating trait, with both phenotypes having shared genetic and environmental components.

In the context of whole genome analyses of human complex traits, there is currently no approach that can fit GCCI effects to disentangle interaction from correlation at the genome-wide level. Yet, ignoring either the genotype-covariate (G-C) correlation or the G-C interaction may cause biased estimates of variance components which form the basis of SNP-heritability or interaction estimation^8^. Random regression-genomic restricted maximum likelihood (RR-GREML)^12; 13^ and G-C interaction (GCI)-GREML^14^ have been used to detect and estimate G-C interaction at the whole genome level for BMI modulated by smoking quantity^12^. However, the analytical approach used in their study was based on univariate models which did not account for G-C correlation, thereby assuming that there is no correlation between the covariates and the outcomes. This can inflate signals indicating the presence of G-C interaction and lead to biased estimates by the failure to account for the G-C correlation. A further limitation with the existing methods is that these cannot be applied to continuous covariates without an arbitrary stratification into discrete exposure groups. Importantly, the methods used for the estimation of SNP heritability (such as GREML^13; 14^ which is based on individual level data, or LDSC^15-17^ based on summary statistics) may give biased estimates for genetic and residual (error) variance if the trait of interest is moderated by (unknown) covariates due to failure in adequate capture of the interaction effects. It is currently not possible to use RR-GREML or GCI-GREML to assess such bias especially when using continuous covariates.

In this study, we develop a whole-genome reaction norm model (RNM) that is computationally flexible and powerful when estimating genome-wide G-C interactions for complex traits. We also extend this approach to a whole-genome multivariate RNM (MRNM) framework to capture fully the GCCI effects, jointly modelling pleiotropy and interactions at the genome-wide level. As the proposed methods will be able to more realistically account for the complexity of GCCI effects, we hypothesize that they will lead to a significant reduction in bias and notably improve the estimation of the genetic architecture of complex traits.

## RESULTS

### Overview of methods

We propose an extension of the whole-genome RNM that can estimate G-C interactions, where covariates can be continuous phenotypes of traits correlated with the response. In a simplified form of this model, the response variable (*y*) representing the main trait is modulated by a continuous covariate variable (*c*) as

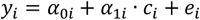

where *α*_0*i*_ and *α*_1*i*_ are the first and second order terms from a random regression of y on c (i.e. the regression coefficients may vary between individuals) and *c_i_* and *e_i_* are the covariate value and the residual effect for the i^th^ individual record (see Methods for the formal model specification and covariance structure). We assessed the power and accuracy in the estimation of G-C interaction for a complex trait modulated by continuous covariates, and compared the performance of RNM with current methods^12-14^ including RR-GREML and GCI-GREML which require stratification of the data into discrete groups due to inability to allow for continuous covariates^12^. In addition, we applied the standard GREML^13; 14^ and LDSC^15-17^ methods to estimate SNP-heritability for the main response variable (*y*), where *y* is modulated by one or more unknown covariates. With this analysis, we are able to explore the potential bias in results obtained by these methods in the presence of non-negligible G-C interactions.

The RNM described above is used to model GCI effects without accounting for G-C correlation. As briefly explained in the Introduction, the same genetic factors can affect both the covariate trait and the main trait (response variable), and at the same time, the covariate trait phenotypes can directly modify the main trait. For example, both BMI and smoking have non-zero SNP-heritability^18^, there is a direct genetic association between BMI and smoking quantity, and BMI is known to be modulated by smoking. Typically, the covariate itself (here, smoking) is affected by genetic effects and residual error (i.e. *c_i_* = *β_i_* + *ε_i_*), and there can be non-negligible correlations between *α*_0_ and *β*, *α*_1_ and *β*, and *e* and *ε*(for the full covariance structure, see Methods). We used multivariate RNM (MRNM) to take into account the G-C correlation, and we compared its performance with the RNM model by assessing differences in bias, type I error rate and power of identifying G-C interactions using simulated data.

The MRNM can be generalised to account for residual heterogeneity or residual-covariate (R-C) interaction, that is

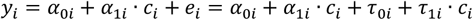

where *α*_0i_, *α*_1i_, and *c*_i_ are defined as above, and *τ*_0*i*_ and *τ*_1*i*_ are the first and second order of random regression coefficients for the residual variance (see Methods).

We compared the performance of previously published methods (RR-GREML^12; 13^ and GCI-GREML^14^) with RNM and MRNM with simulated and real data from the UK Biobank^19^. The models used in these comparisons are summarised in Table S1, a brief description of the UK Biobank and details for the variables used in our analyses are given in Methods and Supplementary Note. In the analyses using the UK Biobank, we modelled BMI as the main trait and fitted separate models using information on pack years of smoking (SMK), neuroticism score (NEU) and the first principal component of genotypes (PC1) as the covariates. Models using PC1 as the covariate were used as a negative control, as the data used in these analyses was stringently restricted according to their ancestry, excluding all participants with values over 6 SD from the mean of the first and second PCs. Due to this stringent restriction based on ancestry, we would expect to see little or no evidence for interaction due to PC1. We used SMK and NEU because of their well-known association with BMI^12; 20; 21^ although the variance and covariance components of the interaction effects were not clearly known.

### Type I error rate, power and estimates in the GCI model

We used simulation (see Methods) to quantify type I error rate and power of detecting G-C interaction for the proposed RNM, RR-GREML and GCI-GREML, without considering G-C correlation. As shown in Figure 1, all methods could control type I error rate under the null model, when there was no G-C correlation and interaction. In contrast, when there were non-negligible G-C interactions, RNM outperformed RR-GREML and GCI-GREML in detecting G-C interactions (Figure 2). The power to detect G-C interaction was slightly higher for RR-GREML compared to GCI-GREML.

**Figure 1.**
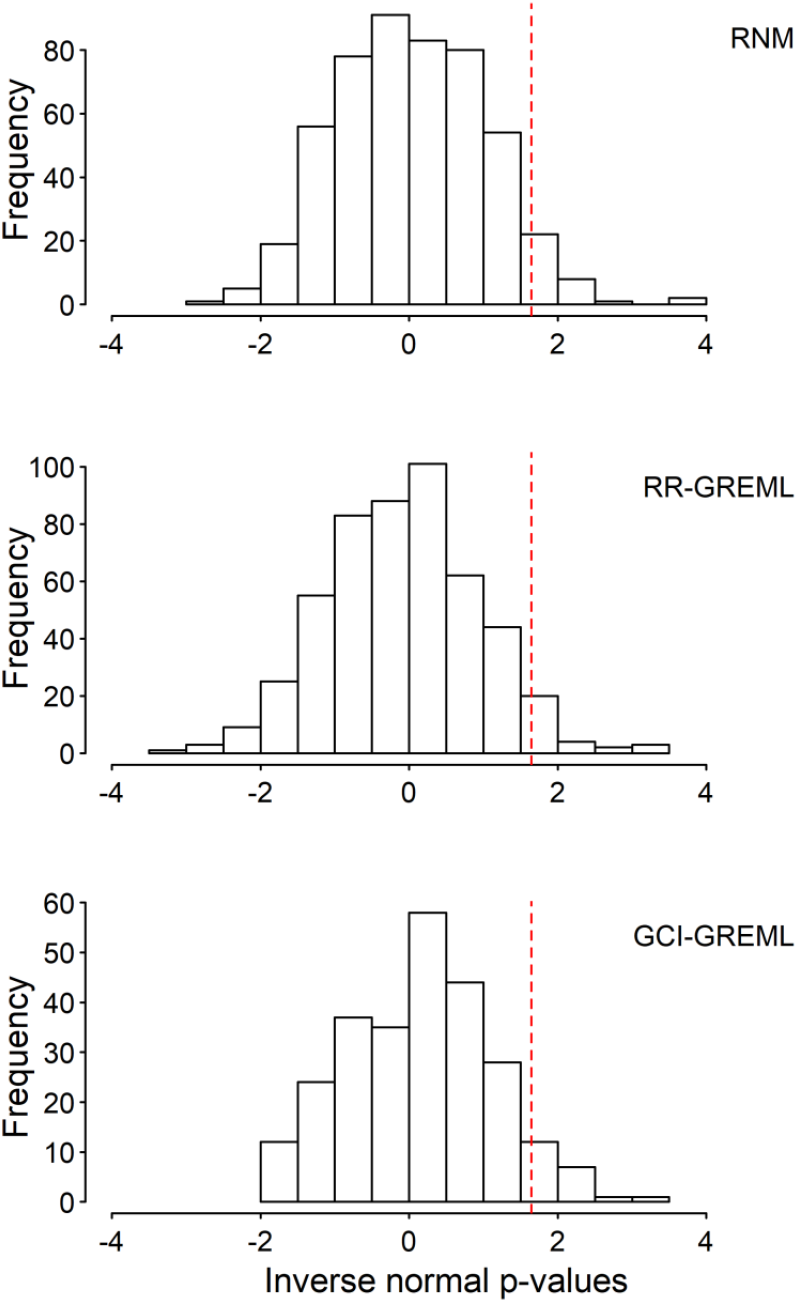
Type I error rate for detecting G-C interaction is under control for RNM, RR-GREML and GCI-GREML. Five hundred replicates of data were simulated under a null model that assumed no genotype-covariate interaction. The model is specified as **y** = **α_0_** + **α_1_**×**c + e** with **c** = **β** + **ε**, where the variance-covariance structure between **α_0_**, **β**, and **α_1_** (in this order) is 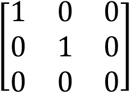 and that between **e** and **ε** is 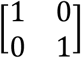. For every replicate, each of the three models was fitted to obtain a p-value for the G-C interaction via a comparison between the null (H_0_) and alternative hypothesis (H_1_) models. For RNM, the H_0_ and H_1_ models were **y** = **α_0_ + e** and **y** = **α_0_** + **α_1_**×**c + e**. For RR-GREML and GCI-GREML, the H_0_ and H_1_ models were **y** = **α_0_ + e** and **y** = **α_0_** + **α_1_**×**c + e**. In RR-GREML and GCI-GREML, samples were arbitrarily stratified into four different groups according to the covariate levels. RR-GREML explicitly estimate residual variance for each of the four groups whereas GCI-GREML assumes homogeneous residual variance across the four groups and estimates a single residual variance. This figure shows the proportions of significant p-values, i.e., type I error rate, for RNM, RR-GREML and GCI-GREML, which are 0.048, 0.048 and 0.034, respectively. Note that p-values are inverse normal transformed, such that the statistical significance level, i.e., 1.65, shown as dashed lines, is equivalent to the 0.05 level before the transformation.

**Figure 2.**
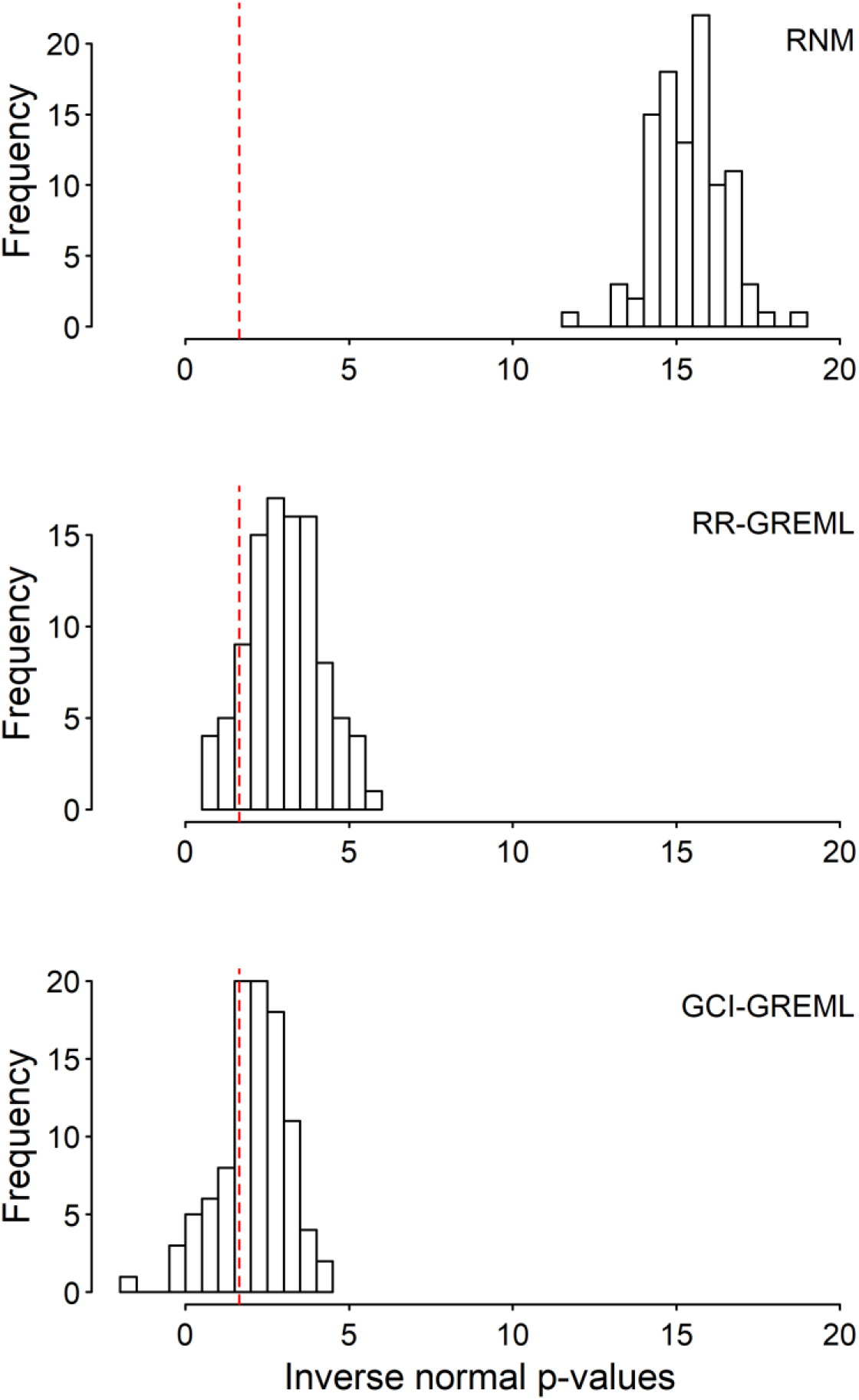
RNM has more statistical power than RR-GREML and GCI-GREML for detecting G-C interaction. One hundred replicates of data were simulated under a model that assumed the presence of a genotype-covariate interaction. The model is specified as **y** = **α_0_** + **α_1_**×**c + e** with **c** = **β** + **ε**, where the variance-covariance structure between **α_0_**, **β**, and **α_1_** (in this order) is 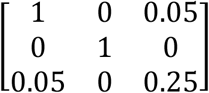 and that between **e** and **ɛ** is 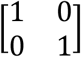. For every replicate, each of the three models was fitted to obtain a p-value for the G-C interaction via a comparison between the null (H_0_) and alternative hypothesis (H_1_) models. For RNM, the H_0_ and H_1_ models were **y** = **α_0_ + e** and **y** = **α_0_** + **α_1_**×**c + e**. For RR-GREML and GCI-GREML, the H_0_ and H_1_ models were **y** = **α_0_ + e** and **y** = **α_0_** + **α_1_**×**c + e**. In RR-GREML and GCI-GREML, samples were arbitrarily stratified into four different groups according to the covariate levels. RR-GREML explicitly estimate residual variance for each of the four groups whereas GCI-GREML assumes homogeneous residual variance across the four groups and estimates a single residual variance. This figure shows the proportions of significant p-values, i.e., statistical power, for RNM, RR-GREML and GCI-GREML, which are 1, 0.9 and 0.69, respectively. Note that p-values are inverse normal transformed, such that the statistical significance level, i.e., 1.65, shown as dashed lines, is equivalent to the 0.05 level before the transformation.

We also tested if the methods can give unbiased estimates for variance components of random regression coefficients underlying the mechanism of G-C interaction. When G-C interactions were present, RNM gave unbiased estimates, whereas estimates from RR-GREML and GCI-GREML differed from true values (Table 1). Note that RR-GREML and GCI-GREML required the stratification of the sample into discrete groups, resulting in an artificial heterogeneity of phenotypic variances across the discrete groups (Figure S1).

**Table 1.**
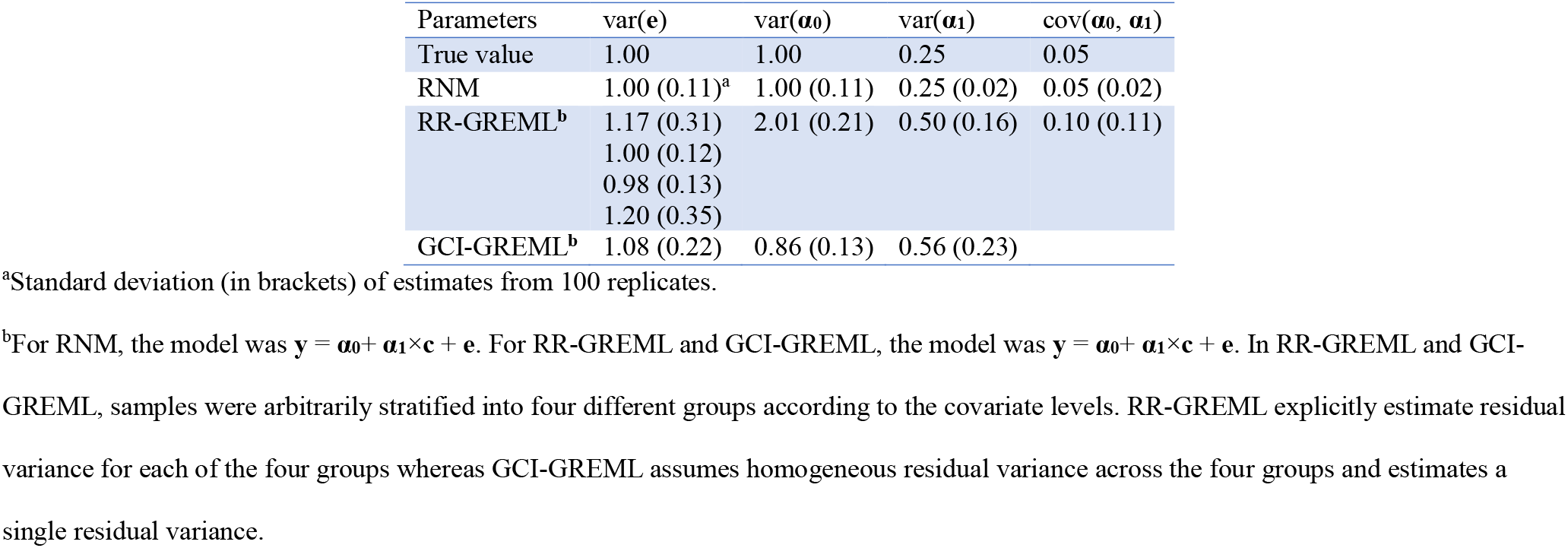
Estimated variance components of random regression coefficients from data simulated under a model that assumed the presence of genotype-covariate interaction.

### Type I error rate, power and estimates in the GCCI model

We also considered the GCCI model in simulations (Methods). As expected, in the presence of non-negligible genetic correlations between the main response and covariate variables (e.g. r≥0.5), we saw spurious signals for G-C interaction in the univariate analysis using the RNM (Figure 3). However, MRNM performed notably better in these analyses, being able to control for type I error rate (0.046) in detecting G-C interaction (Figure 3).

**Figure 3.**
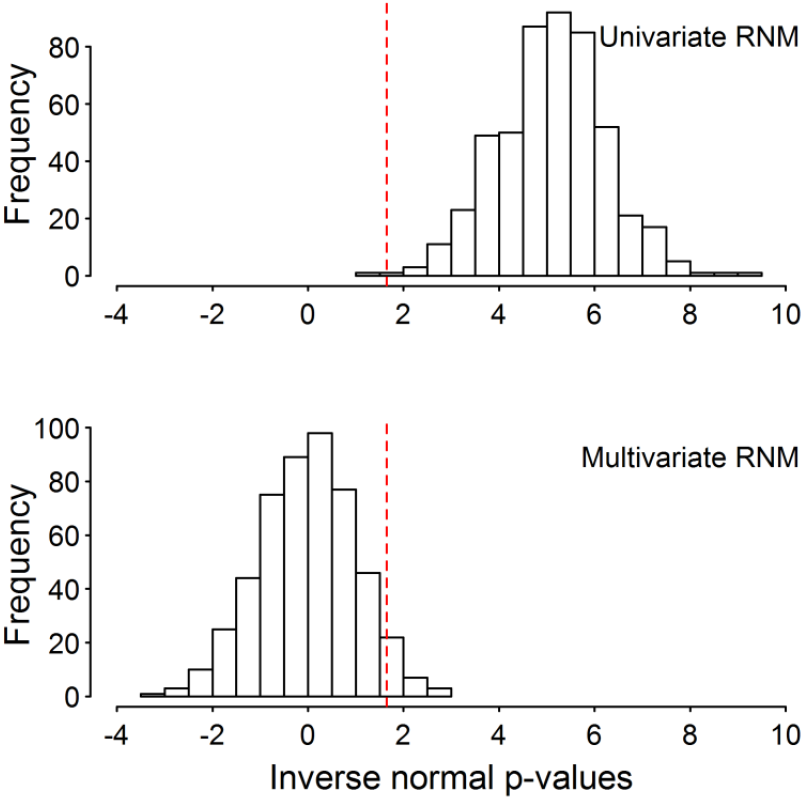
Spurious signals generated by incorrect (univariate) model can be controlled by applying multivariate RNM for detecting G-C interaction. Five hundred replicates of data were simulated under a null model that assumed genotype-covariate correlation but no genotype-covariate interaction. The model is specified as **y** = **α_0_** + **α_1_**×**c + e** with **c**= **β** + **ε**, where the variance-covariance structure between **α_0_**, **β**, and **α_1_** (in this order) is 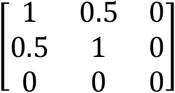 and that between **e** and **ε** is 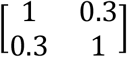. For every replicate, a univariate RNM and a multivariate RNM were fitted separately to obtain a p-value for the G-C interaction by comparing the null (H_0_) and alternative hypothesis (H_1_) model. For the univariate RNM, the H_0_ and H_1_ models were **y** = **α_0_ + e** and **y** = **α_0_** + **α_1_**×**c + e**. For the multivariate RNM, the H_0_ and H_1_ models were **y** = **α_0_ + e** with **c** = **β** + **ε** and **y** = **α_0_** + **α_1_**×**c + e** with **c** = **β** + **ε**. This figure shows the proportions of significant p-values, i.e., type I error rate, for both models, which are 0.998 (univariate RNM) and 0.046 (multivariate RNM). Note that p-values are inverse normal transformed, such that the statistical significance level, i.e., 1.65, shown as dashed lines, is equivalent to the 0.05 level before the transformation.

In the presence of G-C correlations and G-C interactions, both RNM and MRNM performed similarly in detecting G-C interactions (Figure 4) although the significance of the G-C interaction for RNM was slightly inflated due to over-estimated parameters (see var(*α*_1_) for RNM in Table 2). Importantly, MRNM gave unbiased estimates for both G-C correlation and G-C interaction (Table 2).

**Table 2.**
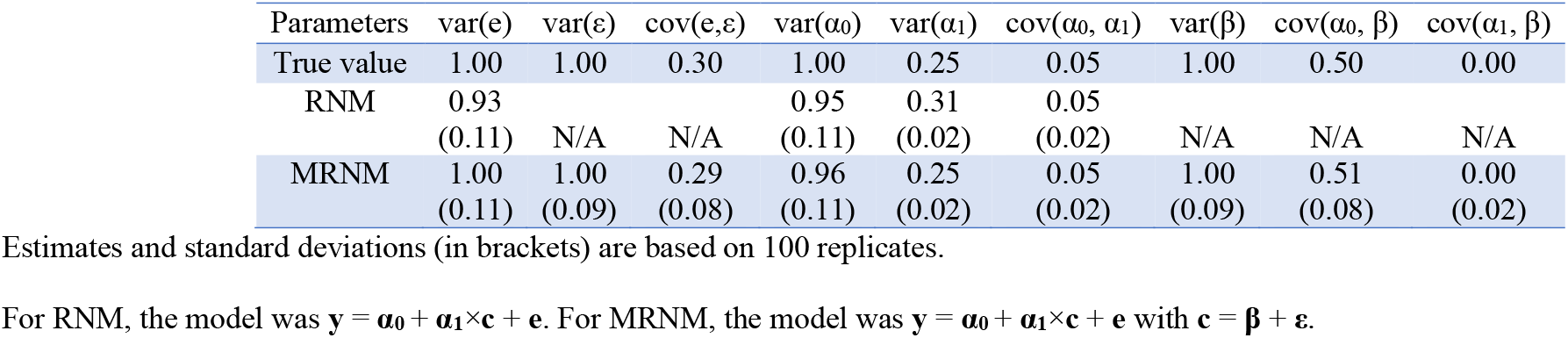
Estimated variance components of random regression coefficients from data simulated under a model that assumes the presence of genotype-covariate correlation (i.e. cov(α_0_, α) > 0) and interaction (i.e. var(α_1)_ > 0). Estimates and standard deviations (in brackets) are based on 100 replicates. For RNM, the model was **y** = **α_0_** + **α_1_**×**c + e**. For MRNM, the model was **y** = **α_0_** + **α_1_**×**c + e** with **c** = **β** + **ε**.

| Parameters | var(e) | var($\varepsilon$) | cov(e,$\varepsilon$) | var($\alpha_0$) | var($\alpha_1$) | cov($\alpha_0, \alpha_1$) | var($\beta$) | cov($\alpha_0, \beta$) | cov($\alpha_1, \beta$) |
| --- | --- | --- | --- | --- | --- | --- | --- | --- | --- |
| True value | 1.00 | 1.00 | 0.30 | 1.00 | 0.25 | 0.05 | 1.00 | 0.50 | 0.00 |
| RNM | 0.93<br>(0.11) | N/A | N/A | 0.95<br>(0.11) | 0.31<br>(0.02) | 0.05<br>(0.02) | N/A | N/A | N/A |
| MRNM | 1.00<br>(0.11) | 1.00<br>(0.09) | 0.29<br>(0.08) | 0.96<br>(0.11) | 0.25<br>(0.02) | 0.05<br>(0.02) | 1.00<br>(0.09) | 0.51<br>(0.08) | 0.00<br>(0.02) |

For RNM, the model was $\mathbf{y} = \alpha_0 + \alpha_1 \times \mathbf{c} + \mathbf{e}$ . For MRNM, the model was $\mathbf{y} = \alpha_0 + \alpha_1 \times \mathbf{c} + \mathbf{e}$ with $\mathbf{c} = \beta + \varepsilon$ .

**Figure 4.**
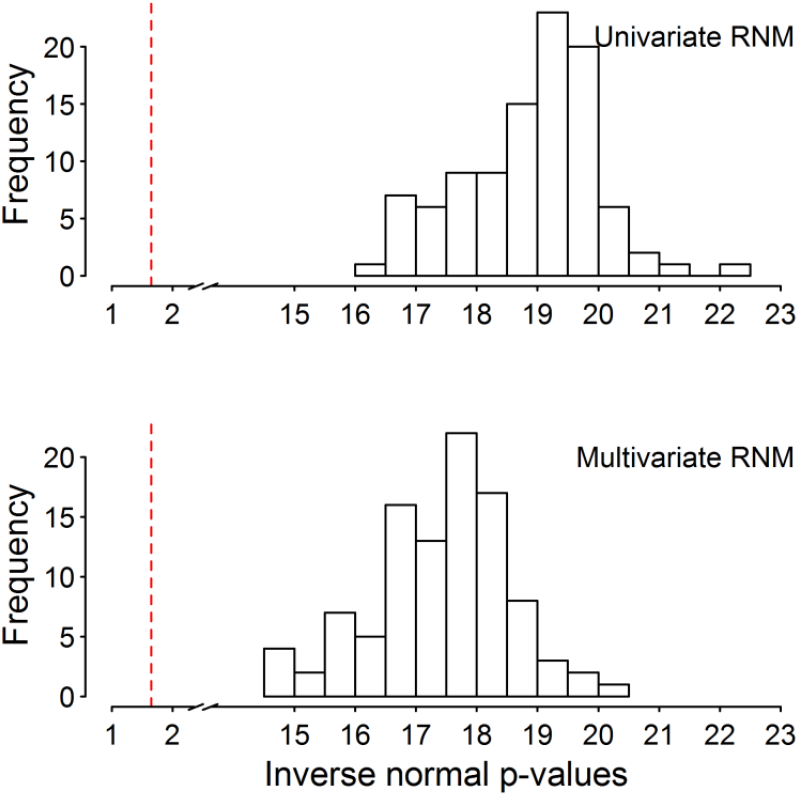
Univariate RNM and multivariate RNM have a similar level of statistical power for detecting G-C interaction. A hundred replicates of data were simulated under a model that assumed the presence of genotype-covariate correlation and interaction. The model is specified as **y** = **α_0_** + **α_1_**×**c + e** with **c** = **β** + **ε**, where the variance-covariance structure of **α_0_**, **β**, and **α_1_** (in this order) is 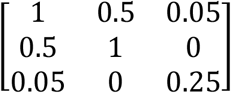 and that of **e** and **ε** is 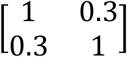. For every replicate, a univariate RNM and a multivariate RNM were fitted separately to obtain a p-value for the G-C interaction by comparing the null (H_0_) and alternative hypothesis (H_1_) model. For the univariate RNM, the H_0_ and H_1_ models were **y** = **α_0_ + e** and **y** = **α_0_** + **α_1_**×**c + e**. For the multivariate RNM, the H_0_ and H_1_ models were **y** = **α_0_ + e** with **c** = **β** + **ε** and **y** = **α_0_** + **α_1_**×**c + e** with **c** = **β** + **ε**. This figure shows the proportions of significant p-values, i.e., statistical power, for the two models, which are 1 for both. Note that p-values are inverse normal transformed, such that the statistical significance level, i.e., 1.65, shown as dashed lines, is equivalent to the 0.05 level before the transformation.

When using RNM, the spurious signals for detecting G-C interaction could be controlled by adjusting the main response for the covariate, i.e. using residuals (as the response) from the regression of the main response on the covariate (Figure S2 and S3). However, such adjustment was crude, and the genuine effects were sometimes over-corrected, again leading to biased estimates especially in the estimated variance of the main effects (Table S2).

### Allowing for residual-covariate (R-C) correlation and interaction

In addition to GCCI, it is possible that the residual effects (ei) are correlated with the covariate (ci) and that there is interaction (RCCI) (see Eq. (3) or Methods). We tested various scenarios for detecting G-C interactions in the presence of R-C correlation and/or interaction (Figures S4–S8). In the absence of G-C interactions but with R-C interactions, type I error rate was well controlled in all methods (Figure S4, Table S3). In the presence of G-C interactions and R-C interactions, RNM had greater power to detect G-C interaction compared to RR-GREML or GCI-GREML (Figure S5 and Table S4). In the absence of G-C interaction but with G-C correlations and RCCI, all three methods were able to control type I errors in detecting G-C interaction (Figure S6 and Table S5). With the full GCCI model in the presence of G-C correlation and interaction, and R-C correlation and interaction, MRNM had greater power than RR-GREML or GCI-GREML (Figure S7 and Table S6). When increasing the variance explained by the G-C interaction, the statistical power reached 100% with all three methods (Figure S8 and Table S7). It is notable that MRNM gave unbiased estimates of the components whereas the other methods generated some degree of bias in the estimation (Tables S5, S6 and S7).

### Bias in the estimated heritability using LDSC or GREML

With simulated data based on the G-C or R-C interaction model, we showed that both the GREML and LDSC overestimated the residual variance for the main response variable hence underestimating the SNP-heritability (Figure 5). When the interaction component explained 10% of the total variance, the estimated residual variance based on GREML or LDSC was 1.5 times higher than the true simulated value (Figure 5). When the variance of the interaction was increased to 25% of the total variance, GREML or LDSC overestimated the residual variance up to 3-fold. However, RNM generated unbiased estimates for the residual variance in most cases. It was noted that the estimated genetic variance was mostly unbiased whether using GREML, LDSC or RNM.

**Figure 5.**
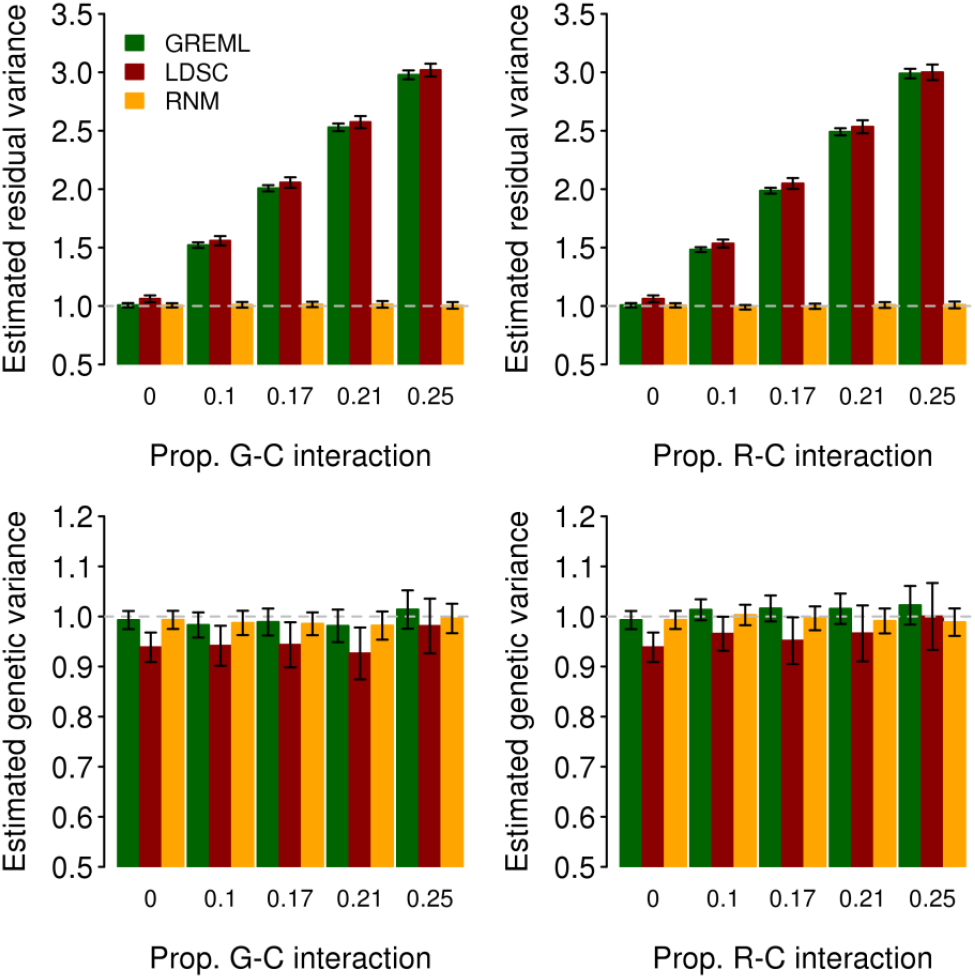
Estimated residual and genetic variances from data simulated under G-C (left) or R-C interaction model (right), using GREML, LDSC and RNM. Prop. G-C or R-C interaction is the proportion of variance due to **α_1_** or **τ_1_ (**see below) in the total phenotypic variance (i.e. var(α**1**))/var(**y**) in G-C interaction model or var(**α_1_**)/var(**y**) in R-C interaction model). Simulation for G-C interaction (**α_1_**): The phenotype data were generated using **y** = **α_0_** + **α_1_**×**c + e** with **c** = **β** + **ε**. The variance-covariance structure of **α_0_**, **β**, and **α_1_** (in this order) is 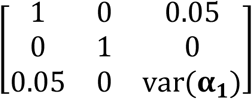 with var(**α_1_**) = 0, 0.25, 0.5, 0.75 and 1 and that for **e** and **ε** is 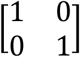. Simulation for R-C interaction (**τ_1_**): The phenotype data were generated using **y** = **α_0_** + **τ_0_** + **τ_1_**×**c** with **c** = **β** + **ε**. The variance-covariance structure of **α_o_** and **β** is 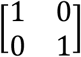 and that of **τ_0_**, **ε**, and **τ_1_** (in this order) is 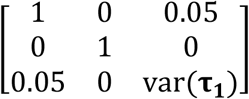 with var(**τ_1_**) = 0, 0.25, 0.5, 0.75 and 1. The error bar is a 95% confidence interval, which was estimated over 100 replicates. The model for GREML is **y** = **α_0_ + e** and the model for RNM in the left panel is **y** = **α_0_** + **α_1_**×**c + e**. The model for RNM in the right panel is **y** = **α_0_** + **τ_0_** + **τ_1_**×**c**.

### Real data

We used the first wave of UK Biobank (UKBB1, see Methods) to compare various models that test interaction using RR-GREML (M1) and GCI-GREML (M2), and the proposed new approaches RNM (M3-7) and MRNM (M8-M12) (Table 3). We conducted GCCI analyses with BMI as the outcome trait using either SMK, NEU or PC1 (negative control) as the covariate of interest.

**Table 3.**
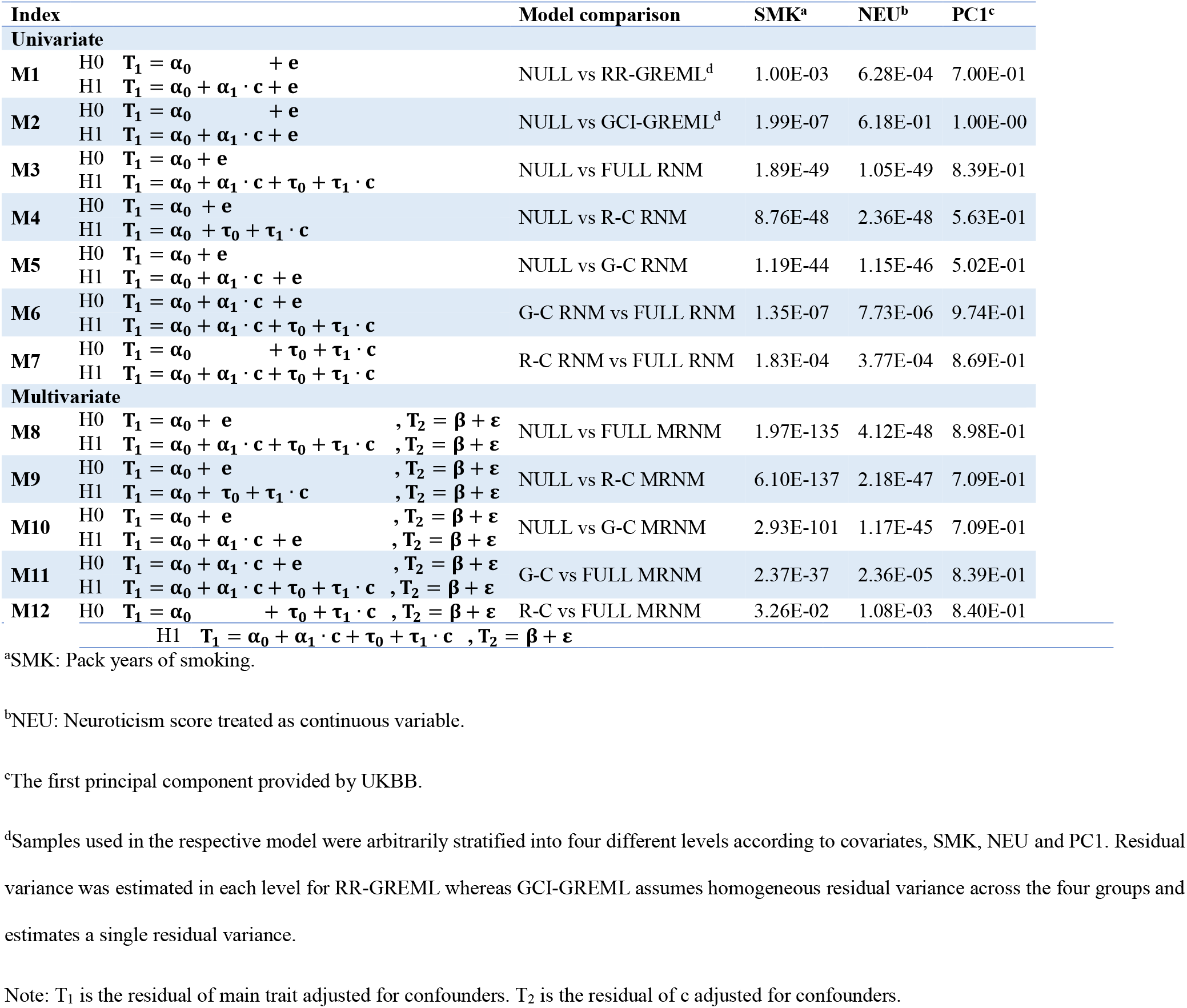
P-values of likelihood ratio tests for model comparisons in UKBB analyses of BMI as the main trait, considering either SMK, NEU or PC1 as a covariate. Note: T_1_ is the residual of main trait adjusted for confounders. T_2_ is the residual of c adjusted for confounders.

Table 3 shows the p-values for interaction effects from the likelihood ratio tests and the corresponding estimates for variance and covariance components are presented in Table S8. We found that BMI was significantly modulated by SMK using RR-GREML (M1, p-value = 1.00E-03) or GCI-GREML (M2, p-value = 1.99E-07), confirming published results^12^. However, these methods did not account for G-C correlation or RCCI (Table S6 and S7). Using RNM (M3–7), we found that the combined G-C and R-C interaction effects were highly significant (M3–5). We then used RNM to test for the G-C interaction corrected for R-C interaction (M7) and found similar results (p-value = 1.83E-04 and var(α_1_) = 0.47 with SE = 0.12) compared to those obtained using RR-GREML (M1) and GCI-GREML (M2). It is noted that residual heterogeneity (reflected by R-C interaction) was partly controlled in M1 and M2 as these models adjusted for group differences with the covariate stratified into four discrete groups, which however generated biased estimates as shown in Table S6 and S7 from simulation. We next applied MRNM to test for interactions, accounting for both G-C interaction and G-C correlation effects (M8–12). We found that the signal for the combined G-C and R-C interaction increased (M8–10) compared to that seen using RNM, which turned out to be mostly due to the increased R-C interaction (M11). It is likely that this was due to the large negative residual correlation between BMI and SMK (Figure 6 and Table S8) which could be more properly modelled in MRNM than in RNM. We finally tested G-C interaction controlled for G-C correlation, and R-C correlation and R-C interaction (M12), and showed that the signal for G-C interaction was negligible (p-value = 3.26E-02) and not significant after considering the number of covariates (n=3) tested in this study. This was probably due to the fact that the non-negligible G-C correlation (Figure 6 and Table S8) would inflate the signal of G-C interaction in M1, 2 and 7 (all based on univariate framework). As shown in the simulations, the MRNM was the most reliable model (Table S6 and S7). Hence, this demonstrates that conclusions from models using the MRNM applied to real data can differ from those obtained using methods based on more simplified models (Figure 6 and Table S8).

**Figure 6.**
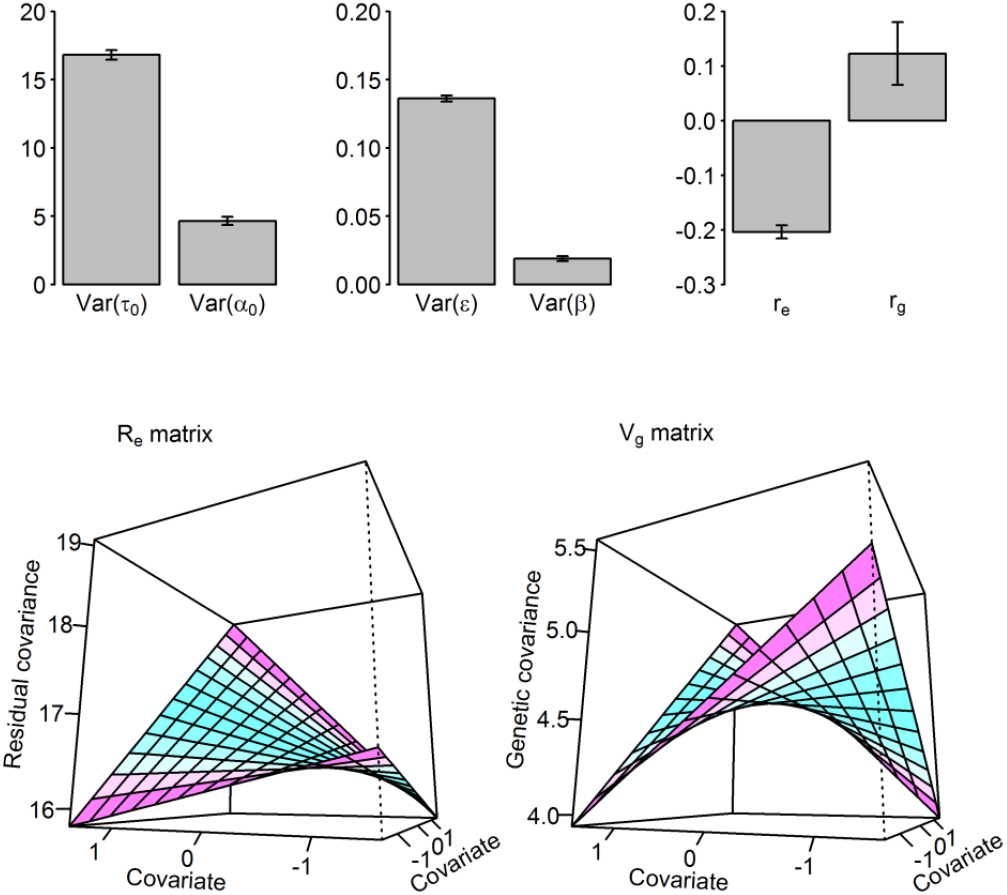
Estimated variance components and correlations from MRNM for BMI-SMK analysis. Var(**τ_0_**): Estimated residual variance for BMI as the main outcome Var(**α_0_**): Estimated genetic variance for BMI as the main outcome Var(**ε**): Estimated residual variance for SMK as the covariate Var(**β**): Estimated genetic variance for SMK as the covariate r_e_: Estimated residual correlation between BMI and SMK r_g_: Estimated genetic correlation between BMI and SMK Error bars are 95% confidence interval R_e_ matrix is the residual (co)variance structure between different covariate levels (see Eq. 4), which is derived using estimated random regression coefficients and polynomial matrix as **R_e_** = **ΦM_y_Φ^′^. Φ** is the matrix of polynomials evaluated at given covariate values, where entries of the first column are all **1**s and the second column is the standardized covariates of respective individuals. **M_y_** is the variance-covariance matrix of estimated random regression coefficients from MRNM as 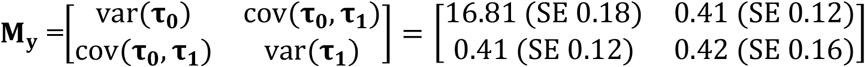. V_g_ matrix in is the genetic (co)variance structure between different covariate levels (see Eq. 2), which is derived based on the estimated random regression coefficients and polynomial matrix as **V_g_** = **ΦM_y_Φ^′^. Φ** is the matrix of polynomials evaluated at given covariate values, where entries of the first column are all **1**s and the second column is the standardized covariates of respective individuals. **K_y_** is the variance-covariance matrix of random regression coefficients estimated from MRNM as 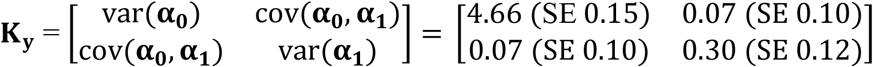.

We also analysed BMI using NEU^20; 21^ as the covariate in the various models, observing evidence for interaction with RR-GREML (p-value = 6.82E-04) but not GCI-GREML(M1 and 2 in Table 3). We found strong G-C and R-C interactions when using either RNM (M3-5) or MRNM (M8-10). Evidence for interaction remained when the G-C interaction effects were adjusted for R-C interaction effects (p-value = 3.77E-04 for M7 and 1.08E-03 for M12) or vice versa (7.73E-06 for M6 and 2.36E-05 for M11). This shows that G-C and R-C interactions are both important and contribute to the shared aetiology between BMI and NEU. As shown in Figure 7, both genetic and residual effects on BMI are significantly modulated by individual NEU while there also is a strong genetic association between them. It was noted that in contrast to BMI-SMK analysis, the results between RNM and MRNM were similar, possibly reflecting different shared genetic and environmental architecture between BMI and NEU, compared to BMI and SMK. The estimated genetic architecture from BMI-NEU analyses is depicted in Figure 7, Table S8 and Figure S9.

**Figure 7.**
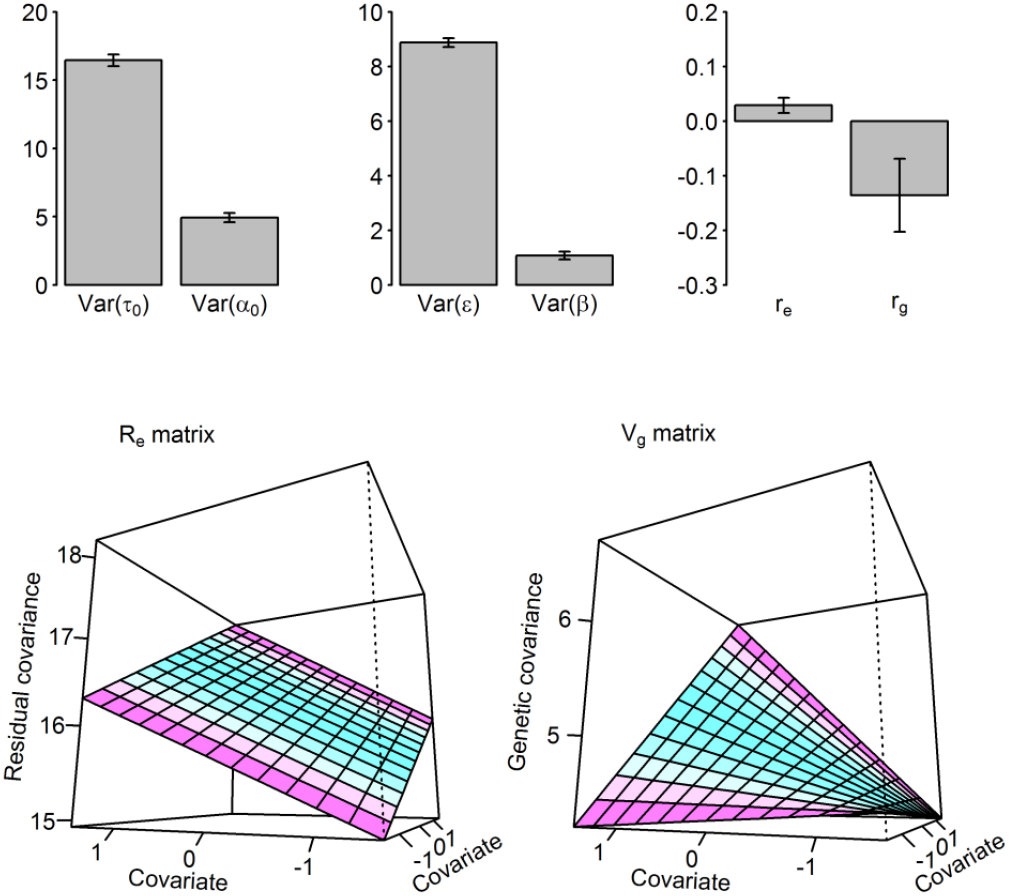
Estimated variance components and correlations from MRNM for BMI-NEU analysis. Var(**τ_0_**): Estimated residual variance for BMI as the main outcome Var(**α_0_**): Estimated genetic variance for BMI as the main outcome Var(**ε**): Estimated residual variance for NEU as the covariate Var(**β**): Estimated genetic variance for NEU as the covariate r_e_: Estimated residual correlation between BMI and NEU r_g_: Estimated genetic correlation between BMI and NEU Error bars are 95% confidence interval Re matrix is the residual (co)variance structure between different covariate levels (see Eq. 4), which is derived based on the estimated random regression coefficients and polynomial matrix as **R_e_** = **ΦM_y_Φ^′^. Φ** is the matrix of polynomials evaluated at given covariate values, where entries of the first column are all **1**s and the second column is the standardized covariates of respective individuals. **M_y_** is the variance-covariance matrix of random regression coefficients estimated from MRNM as 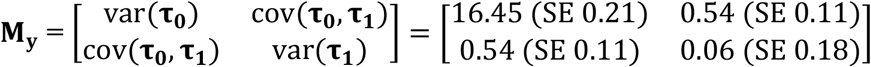. V_g_ matrix in is the genetic (co)variance structure between different covariate levels (see Eq. 2), which is derived based on the estimated random regression coefficients and polynomial matrix as **V_g_** = **ΦM_y_Φ^′^. Φ** is the matrix of polynomials evaluated at given covariate values, where entries of the first column are all **1**s and the second column is the standardized covariates of respective individuals. **K_y_** is the variance-covariance matrix of random regression coefficients estimated from MRNM as 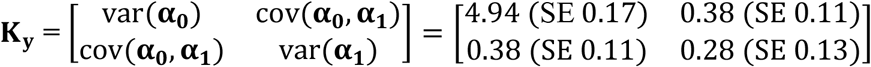.

Lastly, we used PC1 as the covariate in the same analyses (Table 3) and as expected, found no significant interaction effects in this negative control analysis. Compared to SMK or NEU, the R-C interaction was dramatically less (Table 3 and Table S8), probably because PC1 was calculated from genotype data for which the residual component was relatively small. We also found no evidence of G-C interaction, which was probably due to the fact that the sample was so homogeneous such that there was little power to detect interaction effects modulated by population difference.

### An appropriate interpretation of the significance of G-C and R-C interactions for BMI-NEU

The signal of G-C interaction adjusted for R-C interaction (M12) could be underestimated because the R-C model (M10) captured G-C interaction effects and overestimated R-C interaction effects (see the estimated G-C interaction variance component of MRNM R-C model in Table S4-S7). To correct for the bias due to overestimating the R-C interaction, we used the unbiased estimate of R-C interaction variance component from the full model (H1 for M12 in Table 3) that included both G-C and R-C interaction components, on which other estimated parameters were conditioned, when applying the R-C model. We term this the corrected R-C model. P-values from comparing the corrected R-C and the full models would then indicate the significance level of G-C interaction more realistically than p-values from comparing uncorrected R-C and full models (see Figure S10). The same applied to the G-C model, i.e. it overestimated G-C interaction effects, and a corrected G-C model should be used when compared with the full model to get an appropriate significance of R-C interaction effects.

These corrected R-C and G-C models were only applied when the full model was significantly better than either R-C or G-C model such that the estimates from the full model were the most reliable among the models. This was the case for BMI-NEU (the full model had a better-fit than either G-C (p-value = 2.36E-05) or R-C (p-value = 1.08E-03) as shown in Table 3). For BMI-SMK or BMI-PC1, the full model was not significantly better than reduced models after a multiple testing correction. Therefore, we only applied corrected models for BMI-NEU interactions to obtain their corrected significances. In the BMI-NEU analysis, a corrected significance level for G-C and R-C interaction was p-value = 6.98E-10 and 1.54E-13, respectively, which was improved from M11 and 12 in Table 3 because it accounted and corrected for collinearity between two compared model (R-C vs FULL or G-C vs FULL model). As demonstrated in Figure S10, significance level should be carefully interpreted and obtained using the true or a near-true value in the reduced model when testing hypotheses.

### Inflated residual variance using GREML

We observed in simulation data that residual variances for a trait estimated from LDSC or GREML were inflated when there were G-C or R-C interactions (Figure 5 and Table S6 and S6), and this led to underestimates of SNP heritability. Hence, with real data, we tested the differences in the estimates of residual variances for BMI estimated from GREML and RNM (Table 4). For SMK and NEU that had significant interaction effects, the estimated residual variances from GREML were significantly higher than those from RNM (1.89% difference with p-value = 5.99E-04 and 2.04% difference with p-value = 7.12E-03) (Table 4). As expected, there was no significant difference between the models when PC1 was considered as the covariate, because it had no interaction effects. We also fitted both SMK and NEU simultaneously and found that the difference between estimated residual variances from GREML and RNM was increased (3.28% with p-value 1.57E-04) (Table 4). The estimated variance components for the interaction effects from the joint model (Table S9) and the separate models (M4 in Table S8) did not differ. We also observed that the estimated genetic variance varied little between using GREML or RNM (Figure 5 and M3 in Table S3-4), hence biased residual variance directly caused biased SNP-heritability estimates. Inflated residual variance therefore underestimated SNP-heritability, as also observed from an extensive meta-analysis across diverse study-cohorts ^18; 22; 23^ that possibly increased the heterogeneity of covariates shared by the study-samples, hence increased the variance due to G-C and/or R-C interactions (Figure 8). When comparing MRNM and MVGREML, the results did not differ much (Table S10) although there were additional parameters such as cov (α_1_, β) and cov (τ_1_, ε) that were not explicitly parameterised in GREML. We did not fit multiple covariates jointly in MRNM because of our focus on SNP-heritability comparisons based on univariate models (i.e. GREML vs. RNM) and due to the need to control computational demands.

**Figure 8.**
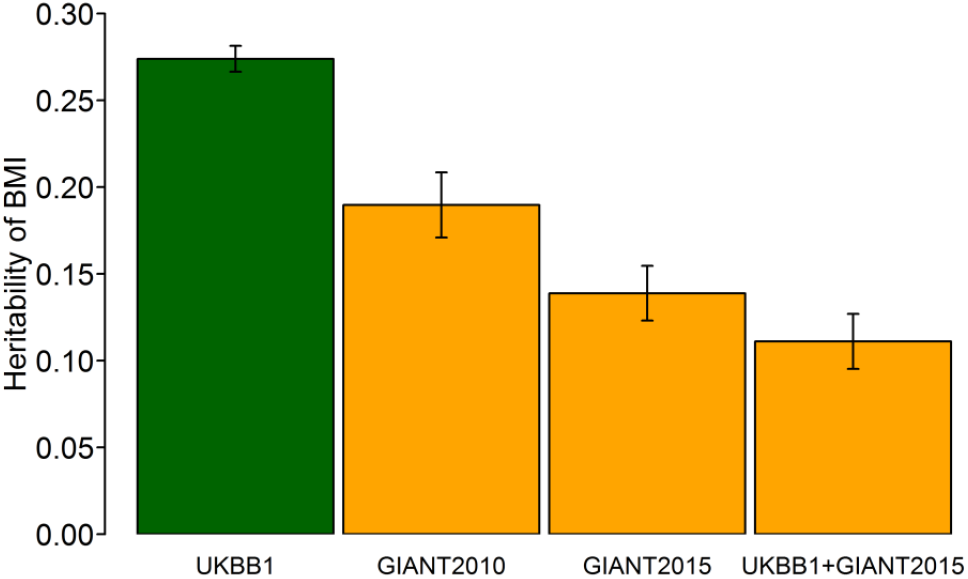
Estimated SNP-heritability of body mass index (BMI) decreases when using increasing numbers of cohorts. The UKBB1 estimate was reported by Ge et al.^18^, which used GWAS summary statistics based on the samples from the first wave of UK Biobank. The GIANT2010 and GIANT 2015 estimates were reported by Duncan et al.^22^, which used GWAS summary statistics based on the GIANT consortium samples from ∼80 and 125 cohorts, respectively. The UKBB1+GIANT2015 estimate was reported by Ni et al.^23^ Bars are 95% confidence interval.

**Table 4.** Statistical tests for the difference between residual variances of BMI estimated from RNM^a^ and GREML^b^.

|  | **Difference<sup>c</sup>** | **SE<sup>d</sup>** | **Difference in %** | **SE of Difference in %** | **h<sup>2</sup> (GREML)** | **h<sup>2</sup> (RNM)** | **P<sup>e</sup>** |
| --- | --- | --- | --- | --- | --- | --- | --- |
| **SMK** | -0.316 | 0.092 | 1.887 | 0.549 | 0.221 | 0.224 | 5.99E-04 |
| **NEU** | -0.336 | 0.125 | 2.044 | 0.760 | 0.227 | 0.231 | 7.12E-03 |
| **PC1** | 0.016 | 0.088 | -0.095 | 0.523 | 0.227 | 0.227 | 8.56E-01 |
| **SMK-NEU<sup>f</sup>** | -0.588 | 0.156 | 3.279 | 0.870 | 0.227 | 0.233 | 1.57E-04 |

<sup>a</sup>Alternative model (H1) of M3 in Table 3.
<sup>b</sup>Null model (H0) of M3 in Table 3.
<sup>c</sup>Difference = the residual variance estimated from RNM – the residual variance estimated from univariate GREML.
<sup>d</sup>Standard error of the difference was calculated based on the theory in Supplementary Note.
<sup>e</sup>P value was obtained based a two-tailed Wald test using the difference of residual variances and its SE.
<sup>f</sup>The model jointly fitted both SMK and NEU as multiple covariates (see Methods).

### Meta-analysis approach and validation

For very large datasets, our proposed approach may become computationally infeasible. A solution could be to divide the data in various subsets and undertake a meta-analysis. In this section, we show that a meta-analysis^24^ of GCCI and RCCI results across difference data subsets is useful and reliable. We simulated phenotypes using UKBB1 genotype data and compared results from meta-analysis of multiple sub-samples with results from each individual sub-sample.

As expected, the values of –log10(P) and likelihood ratio in the meta-analyses were larger than those in each single study (Figure S11). The power increased further as the number of studies (and the total sample size) increased as shown in Figure S11. The correlation between p-values from a meta-analysis based on two groups and p-values from data combining two groups approached to one when the sample size in each group increased to 10K although the regression slope was less than one (Figure S12). As expected, with the same sample size, the power of meta-analyses decreased with the number of groups (e.g. 10K x 2 vs. 4K x 5 in Figure S12) although it was still higher than that from a single group (Figure S11). This indicates that our approach combined with meta-analysis can be applied to any sample size, ensuring that the power keeps increasing with further additions to large-scale biobank data. The increased power in meta-analyses was also evident in real data analyses. We randomly divided the UKBB1 data set into two groups of equal size (33,140 each for SMK, 27,179 each for NEU and PC1) and obtained meta-analysed p-values (Table S11) and estimates (Table S12). In agreement with the simulation, the meta-analysed p-values were not substantially different from those based on the whole data that greatly improved the power compared to using a single study (g1 or g2) (Table S11).

We further performed meta-analyses across the UKBB1 and the second wave of UK Biobank data (UKBB2). UKBB2 excluded the overlapping and highly related samples from UKBB1, and it was used as an independent validation data sets (see Methods for more detail). From the meta-analyses, the significance of R-C interaction effects for BMI-SMK and BMI-NEU increased from P-value = 1.97E-135 to P-value <0E-300 and from 4.12e-48 to 2.73E-121, respectively (M8 in Table S13). G-C interaction effects for BMI-NEU became more significant and the p-values decreased from 1.1E-03 to 5.7E-05 (M12 in Table S13). The meta-analysed estimated variance components are shown in Table S14.

##### Box 1. Summary

1. For continuous covariates, the proposed RNM is a more appropriate model, compared to RR-GREML and GCI-GREML.
2. Covariates can be regulated by genetic and environmental factors that are possibly shared with the main response (GCCI and RCCI effects), which is the most plausible mechanism for many complex traits. It is desirable to model GCCI and RCCI effects appropriately (using multivariate RNM).
3. LDSC and GREML estimates for SNP-heritability should be carefully interpreted or revisited if covariate information is available (Figure 8), and (M)RNM can access the biasedness (Table 4).
4. The proposed models can be applied to any sample size including from large-scale biobank data by meta-analysis of results from sub-samples, for which the analyses are computationally feasible.
5. Our proposed approach is flexible and allows for multiple covariates to be fitted simultaneously (Table 4).

## DISCUSSION

Complex traits are determined by both genetic and environmental effects. Some environmental covariates of complex traits may themselves be determined by genetic and non-genetic factors. Genotype-covariate correlation and interactions (GCCI) and residual-covariate correlation and interaction effects (RCCI) may be important underlying factors shaping complex trait phenotypes^25^, yet not many studies have conducted analyses to detect these effects jointly in one model because of a lack of proper analysis models. In this study, we propose a flexible (multivariate) RNM to estimate genotype-covariate correlation and interactions and residual-covariate correlation and interaction effects for complex traits, which is powerful and reliable. A further benefit with our proposed model is that it can fit continuous covariates that must otherwise be stratified into discrete groups if using the currently available methods of RR-^12; 13^ or GCI-GREML^14^.

We showed in simulations that current univariate approaches including RR-^12; 13^ and GCI-GREML^14^ gave biased estimates (Table S5, S6, and S7) and lower power (Table S6 and S7 or Figure S5, S7, and S8) in general, compared to the proposed (multivariate) RNM. When G-C correlations were ignored, there were spurious or inflated signals for G-C interactions and estimations were biased. Although using adjusted phenotypes corrected for the covariate (regressing phenotypes on the covariate) could control false positives, the estimates could be significantly biased (Table S2). In contrast, the proposed multivariate RNM could effectively control false positives while providing unbiased estimates and reasonable power. This may have a significant implication in obtaining unbiased estimation of the genetic architecture of complex traits. In line with simulations, the results from real data analyses (Table 4) of BMI with pack years of smoking and neuroticism as covariates also showed that the signals for significant G-C interaction might be biased in M1, M2 and M7.

The methods currently used for estimation of G-C interactions, the RR- and GCI-GREML methods, require that for a continuous covariate (e.g. SMK), the outcome of interest (e.g. BMI) should be stratified into multiple discrete groups according to the level of the covariate. This causes heterogeneous phenotypic variance across the groups (Figure S1), which may have non-negligible effects on the estimation of genetic and interaction components in the methods (as shown in Tables S6 and S7). Moreover, the discrete grouping ignores the difference of covariate values for the individuals within each group, and results in some loss of information. In contrast, RNM or MRNM is a flexible model to fit a continuous covariate and gives unbiased estimates in the simulation (Table S6 and S7). With real data, the estimated interaction components for the BMI-SMK analysis were 0.47 (SE=0.14), 1.15 (SE=0.23) and 0.30 (SE=0.12) for RR-GREML, GCI-GREML and MRNM, respectively. The difference between the methods could be explained by the reasons above, i.e. arbitrary grouping and confounding interaction/correlation for the RR- and GCI-GREML methods. Based on the analysis method that we believe to be the most appropriate for the data (MRNM) the G-C interaction estimate was much reduced and only borderline significant while R-C interaction was much more significant.

In the presence of G-C or R-C interactions, estimated SNP-heritability of the main response variable by GREML or LDSC could be biased (Figure 5). The biased estimates reflect that the interaction effects are absorbed by residual variance and the overall estimated residual variance was inflated. The residual variance estimated from GREML was significantly higher than that from RNM for the BMI-SMK, BMI-NEU or BMI-SMK/NEU analysis using the real data (Table 4). Currently reported SNP-heritabilites estimated based on meta analysis of GWAS summary statistics from diverse study-cohorts tend to be lower when the number of study-cohorts is larger (as a proxy of heterogeneity) (Figure 8), which can be partly explained by not properly modelling G-C and R-C interactions. This observation has an important implication because estimates from such meta-analyses (using LDSC) should be carefully interpreted when known key covariates were not included in the GWAS analysis model that generated the input for the LDSC analysis.

In this study, we found a strong negative R-C correlation (−0.20) and weak positive G-C correlation (0.12) between BMI and smoking, which may support the phenomenon observed in several studies that heavier smokers tend to have lower BMI. The R-C interaction was shown to be highly significant (p-value = 6.1E-137 (M9) in Table 3). This suggests that the information about R-C interaction component is crucial such that that the main phenotypes (BMI) can be possibly controlled by changing the covariate (SMK), provided that the covariate is modifiable. In this example, the implication is that the intervention of increasing smoking could be used to control BMI^26^. While in this example the advice may not be practical for other health reasons, the principle can be used to other traits and diseases with modifiable covariates. The information from the G-C correlation and interaction can be useful for an early intervention (e.g. genomic medicine) although the magnitude of the effects are relatively small, compared to R-C components. We also investigated NEU and found strong G-C and R-C interactions (p-value = 4.12E-48 (M8)), indicating the personality trait NEU is a major covariate influencing the environmental factor for BMI as well as revealing a novel genetic architecture of BMI to interact with different levels of NEU (Figure 7). Both genetic and residual variances of BMI are significantly modulated by NEU, as well as there is significant genetic correlation between BMI and NEU. We included analyses using PC1 as the covariate in the model, conducting this analysis was as a negative control analysis (i.e. little variance among the homogenous sample), and, as expected, there were no significant interactions (Table 3). In other circumstances, for example when using diverse samples from the population or even across different ethnic populations, then analyses that fit PC1 as a covariate might generate significant interaction estimates.

The proposed approach can be extended to include epigenetic and gene-expression data as novel covariates, which could reveal the genetic architecture associated with interactions and correlations that involve such variables. The proposed approach will enable us to make a better use of the extensive data resources available, with the prospect of transforming our ability to gain biological insights from genetic, epigenetic and gene-expression data. It is possible to use models that fit multiple covariates simultaneously as we did for fitting both SMK and NEU jointly (Table 4), which increase the proportion of the total phenotypic variance explained by the interaction components. Genomic partitioning analyses to describe GCCI and RCCI effects across the genome will be also useful to shed light on the latent genetic architecture of complex traits and diseases, which is possible by using the proposed approaches in this study.

We did not perform an inverse normal transformation (INT) for the pre-adjusted phenotype (e.g. BMI). In contrast, such transformations were vigorously applied in the analysis Robinson et al.^12^. They also applied INT within each group after the arbitrary stratification according to the level of the covariate, which is a stringent adjustment. In this study, however, we did not use INT because 1) the distribution of phenotypes (BMI) was reasonably normal, 2) using INT is not always the best solution because it may bias the significance and estimation^27^ 3) phenotypes were already adjusted for the covariate and 4) there was negligible difference with and without INT for the whole phenotypes in our analyses. As shown in Figure S1, the arbitrary stratification could generate artefact heterogeneity for phenotypic variance between groups, for which it may not be feasible to correctly control such artefact heterogeneity and generate unbiased estimates. Nonetheless, discrete grouping causing this inconsistency is not applied to RNM or MRNM.

An alternative approach to disentangle interaction from association is through the classical structural equation models^8^ applied to twin- or pedigree-based data. However, availability of such data is limited, restricting our ability to study GCCI effects for a wide range of complex traits and covariates. For example, phenotypes moderated by ancestry components (e.g. ethnic composition in humans or breed composition in animals) cannot be studied by an approach that is based on twins or relatives. It is also difficult to disentangle the genetic and shared environmental effects when using a pedigree-based approach. Standard REML packages (e.g. ASReml^28^) can be used to test the GCCI effects although it is questionable that the classical REML algorithm, which has been optimised for pedigree-based studies, can be computationally tractable when fitting genetic covariance structures based on genomic information^13^. Therefore, it maybe infeasible investigate the GCCI effects using the classical REML packages although they have been applied widely in livestock^29; 30^ and ecological genetics^9; 31; 32^ to explore the phenotype-genotype relationship across environmental gradients. When extending analyses to cover large-scale data such as the UK Biobank^19^, it is essential to develop computationally efficient methods that also correctly capture the GCCI effects, based on genomic information.

There are a number of limitations in this study. Firstly, we did not consider higher order interactions, considering only interaction of order = 2 for both G-C and R-C interactions. A further study is required to validate performance with higher orders interactions to generalise the proposed approach. Secondly, our approaches are flexible, but computationally demanding. For a large data set, it may not be computationally feasible to conduct an analysis within a short time although the meta-analysis approach can get around the problem. Thirdly, the proposed methods do not estimate the direction of causality that can be determined by existing methods, e.g. Mendelian Randomization. However, it is desirable to develop an efficient approach to determine the direction of causality in the context of MRNM (based on random effects models). Fourth, in application to real data we do not take account of ascertainment biases that may generate interactions and correlations in the sample of data which means that our results may not be representative of the populations from which the samples are drawn^33; 34^ although linear mixed models, on which RNM and MRNM are based, are robust to such bias^35; 36^.

In conclusion, we showed that the multivariate RNM is able to effectively disentangle interaction from correlation and to generate unbiased estimates for G-C and R-C components. The concept of GCCI and RCCI is more plausible in explaining the genetic architecture of complex traits associated/interacted with covariates, which will shift the paradigm from a univariate to multivariate framework and from linear to non-linear models in complex trait analyses.

## METHODS

### Reaction norm model (RNM)

To account for phenotypic plasticity and norms of reaction in response to different covariate or environmental conditions among samples^29; 30^, the dependent variable for individual *i* can be modelled as

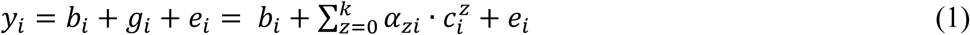

where *y_i_* is the phenotypic observation, *b_i_* represents fixed effects such as sex, *g_i_* is the random genetic effect, *α_zi_* is the *z*th order of random regression coefficients (*z* = 0 ~ *k*), *c*_i_ is the covariate value, and *e_i_* is the residual effect. The variance-covariance matrix of random regression coefficients (**K**) is

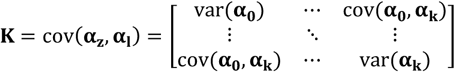

and the genetic (co)variance between *N* individuals, each with a covariate value, is

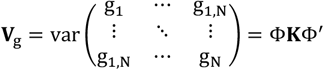

where **Φ** is the matrix of polynomials evaluated at given covariate values. Given that this model does not explicitly parameterise the correlation between *y_i_* and *c*_i_, it naively assumes that *y_i_* and *c*_i_ are uncorrelated. For this reason, this model is also referred to as a genotype-covariate interaction (GCI) model.

### Multivariate reaction norm model (MRNM)

The naïve assumption of the univariate RNM (or GCI model) that *y_i_* and *c*_i_ are uncorrelated is often violated. In a more proper model, the covariate for individual *i* is decomposed as *c_i_* = *μ_i_* + *β_i_* + *ε_i_* where *μ_i_* is fixed effects (e.g. sex), *β_i_* is the random genetic effect, and *ε_i_* is the residual effect. When considering the main response (*y*) and covariate (*c*) together, the covariance structure of the multivariate genetic components becomes

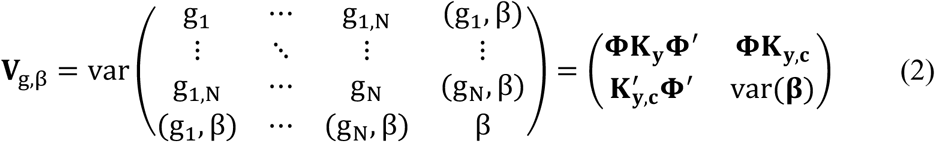

where **K**_*y*_ is the same as **K** defined above, and **K**_*y,c*_ is the covariance matrix of multivariate random regression coefficients, defined as

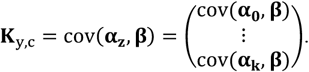

The covariance structure of the multivariate residual components is

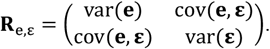

Under this model, the variance-covariance structure consists of var(**α_0_**), var(**α_k_**), cov(**α_0_**, **α_k_**), var(**e**), var(**β**), var(**ε**), cov(**α_0_, β**), cov(**α_0_, β**), and cov(**e, ε**). For this reason, this model is referred to as a genotype-covariate correlation and interaction (GCCI) model. Importantly, values for cov(**α_0_, β**), cov(**α_k_, β**) or cov(**e, ε**) are often non-negligible. Neglecting these terms can cause confounding between G-C correlation and interaction, thereby generating spurious signals and biased estimates for the interaction. Yet many studies do not account for G-C correlations when estimating and testing G-C interaction^12^.

### Multivariate reaction norm model (MRNM) accounting for heterogeneous residual variance, i.e. residual-covariate correlation and interaction (RCCI)

The models we described so far assume that the residual variance, var(**e**), is homogeneous across different values of the covariate. However, it is often possible that residual-covariate (R-C) correlation and interaction exist, resulting in heterogeneous residual variance. To account for this possibility, MRNM can be further extended as

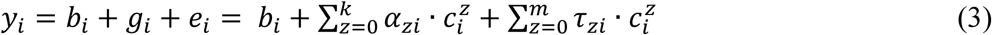

where the residual term in model (1) is expressed as the random regression coefficients *τ_zi_* with a function of the zth order polynomial of the covariate (*z* = 0 ~ *m*).

The variance-covariance structure of the genetic effect for this model is the same as for the multivariate reaction norm model described in Eq. (2) above. The variance-covariance structure of the multivariate residual components becomes

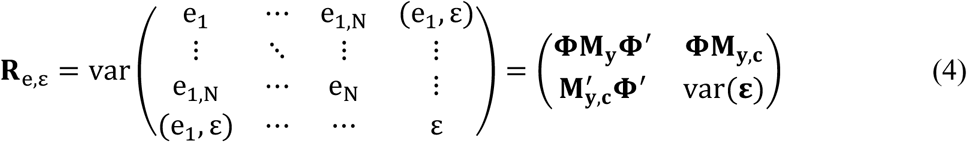

where **M_*y*_** is the variance and covariance matrix of random regression coefficients for the residual components and can be written as

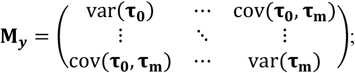

**M_*y,c*_** is the covariance matrix of multivariate random regression coefficients for the residual components and can be expressed as 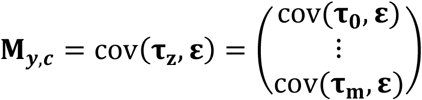 and *var*(ε) is the residual variance of the covariate.

### RNM with multiple covariates

RNM can be further extended to include more than one covariates. The main trait for individual *i*, *y_i_*, can then be expressed as

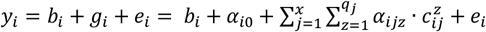

where *x* is the number of random effects, which equates to the number of covariates included. Each random effect has random regression coefficients, *α_ijz_*, with z = 1 ~ *q_j_*, fitted with the j^th^ covariate. It is assumed that there is no correlation between the random effects.

The variance-covariance matrix for each random effect is the same **K** as above, that is of random regression coefficients (**K**) is

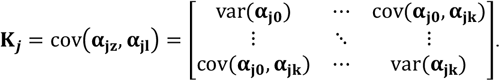

Similarly, the genetic (co)variance between N individuals, each with a set of covariate values, can be obtained as 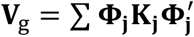 where **Φ_j_** is the matrix of polynomials evaluated at given values of the jth covariate.

This can be feasibly extended to MRNM with RCCI although the number of parameters increases exponentially. All models described above (i.e., RNM and MNRM) can be fitted using MTG2^13^.

### Simulated data

Phenotypic simulation was based on individual genotypes from the GWAS data of the Atherosclerosis Risk in Communities Study (ARIC) cohort. We used autosomes only and applied the standard quality control (QC) to genotypes, which included MAF > 0.01, SNP call rate > 0.95, sample call rate > 0.95 and Hardy-Weinberg Equilibrium p-value > 0.001, keeping qualified genotyped SNPs. After the QC, 583,058 SNPs and 8,291 individuals remained. We estimated pair-wise relatedness from the remaining SNPs and randomly excluded one individual from each pair with an estimated relatedness greater than 0.05. This reduced the sample to 7,263 individuals.

#### Simulation under GCCI model

We simulated phenotypes for the main response (*y*) and covariate (*c*) under the GCCI model with the first order interaction effect, i.e. *k*=1. In the simulation, we used the following covariance structure for the **K**_y_ matrix in Eq. (2)

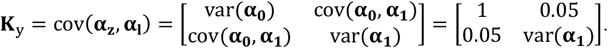

We used a wide range of the G-C interaction with *var*(***α*_1_**) set at 0, 0.25, 0.5, 0.75 or 1. For the covariate, *c_i_* = *μ_i_* + *β_i_* + *ε_i_*, var(**β**) and var(**ϵ**) were set at 1.

The **K**_y,c_ matrix (Eq. 2) was specified as

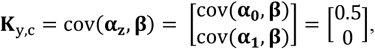

The values for the **R**_e,ε_ matrix were used in the simulation as

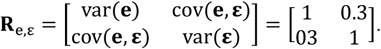

For null model, we set the interaction variance, i.e., var(alpha1), at zero. We assessed type I error rate and power of detecting G-C interaction under the null and full model, respectively.

#### Simulation under GCCI model with R-C correlation and/or interaction

Similar to the simulations above, we used values for the **K**_y_ matrix in Eq. (2) as

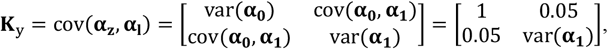

We performed simulations with var(*α*_1_) set at 0.25 or 1. For the covariate, both *var*(***β***) and *var*(***ϵ***) were set at 1.

The **K**_y,c_, ***M_y_*** and ***M_y,c_*** matrix (Eq. 3 and Eq. 4) were specified as follows:

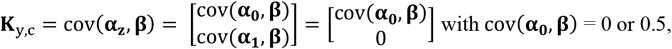

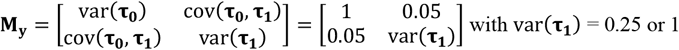

and

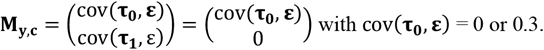

We assessed bias, type I error rate and power for LDSC ^15; 16^, GREML^14^, MVGREML^13^, RR-GREML^12^, GCI-GREML^14^, RNM, and MRNM using simulated data generated as above. Subsequently, we compared the performance of the methods.

### Real data

#### Data and Quality control

We used the UK Biobank data^19^, which initially contained 488,377 individuals and 92,693,895 imputed SNPs across autosomes. Stringent quality control was applied to the genotype data at both individual and SNP levels. Specifically, we excluded individuals who met one of the following criteria: 1) does not have white British ancestry, 2) has a genotype missing rate > 0.05, 3) whose reported gender does not match with the gender inferred using genotype data, and 4) has a putative sex chromosome aneuploidy. At the SNP level, we excluded SNPs with an INFO score < 0.6, with a minor allele frequency (MAF) < 0.01, with a Hardy-Weinberg equilibrium P-value <1E-4, or with a call rate < 0.95. For multiple records of the same SNP, we randomly selected one and removed duplicates. We also excluded ambiguous SNPs and only kept HapMap 3 SNPs.

In addition, we excluded individual population outliers, namely individuals with the first or second PC outside six standard deviations of its respective population mean. For individuals who were in both UKBB1 and UKBB2, we calculated the discordance rate between imputed genotype of the two versions for each individual and for each SNP, and excluded individuals and SNPs with a discordance rate lager than 0.05. We also excluded one individual randomly from any pair with a genomic relationship larger than 0.05. After the QC above, 288,866 individuals and 1,130,918 SNPs remained. Of these remaining individuals 91,472 were from UKBB1, who were used in the main analyses and 197,394 were from UKBB2, who were used in the validation and meta-analyses.

#### Main response variable and covariates

We applied the novel (M)RNM model using BMI as the main response variable to estimate the GCCI/RCCI components with each of several covariates, including pack years of smoking (SMK), neuroticism score (NEU) or the first principal component (PC1) provided by the UK Biobank. We also fitted the model that includes multiple covariates (e.g. SMK and NEU) jointly, i.e., RNM with multiple covariates. For all analyses, covariates were standardized as mean zero and variance 1. Prior to model fitting, we adjusted the main response variable (BMI) for confounders including genotype batch, assessment centre at which participant consented, year of birth, sex, age, diet variation, diet change, the first 15 PCs, SMK, weekly alcohol consumption (ALC) and Townsend deprivation index at recruitment (TDI). The distribution of each covariate is in Figure S13.

When including the covariate (i.e. SMK, NEU, or PC1) as the second trait in a MRNM, it was also pre-adjusted for the confounders in a similar way as for the main trait (i.e., BMI). For instance, as the second trait in a MRNM, SMK was pre-adjusted for BMI, genotype batch, assessment centre at which participant consented, year of birth, sex, age, diet variation, diet change, the first 15 PCs, ALC, and TDI. NEU was pre-adjusted for BMI, genotype batch, assessment centre at which participant consented, year of birth, sex, age, diet variation, diet change, the first 15 PCs, ALC, TDI and SMK. PC1 was pre-adjusted for BMI, genotype batch, assessment centre at which participant consented, year of birth, sex, age, diet variation, diet change, the first 15 PCs except the first one (PC1), ALC, TDI, SMK and NEU.

Detailed information regarding covariates used in the interaction models is described below and that for other confounders used to adjust the main phenotypes is in Supplementary note.

#### SMK

We combined pack years adult smoking as proportion of life span exposed to smoking (UK Biobank data field: 20162) and ever smoked (UK Biobank data field 20160) as SMK. The distribution of SMK is in Figure S13. For RR-GREML and GCI-GREML, following Robinson et. al.^12^, we stratified SMK into four levels: 8,773 individuals with SMK > 0.8, 9,192 individuals with 0.5 ≤ SMK ≤ 0.8, 11,741 individuals with 0< SMK <0.5, and 36,575 individuals with SMK =0 (i.e. never smoked).

#### Neuroticism score (NEU)

The neuroticism score (data field 20127) of a given individual was indexed by the number of ‘yes’s to 12 touchscreen questions that evaluate neurotic behaviours. The distribution of NEU is in Figure S13. For RR-GREML and GCI-GREML, we stratified the data into four groups according to NEU level: 20,901 individuals with NEU ≤ 2, 16,161 individuals with 2 < NEU ≤ 5, 10,895 individuals with 5 < NEU ≤ 8, and 6,417 individuals with 8 < NEU ≤ 12.

#### PCI

PCs were pre-calculated by the UK Biobank. Detailed information regarding the calculation is described elsewhere^37^. Briefly, PCs-loadings were estimated using fastPCA^38^ based on 407,219 unrelated individuals and 147,604 markers that were pruned to minimise linkage disequilibrium, onto where all samples were projected, to generate a set of PC scores. For RR-GREML and GCI-GREML, we stratified the sample into four groups based on quartiles of PC1.

#### Meta analyses of real data

The proposed MRNM requires individual-level genotype data, which makes it computationally demanding. As sample size increases (e.g. the second wave of UK Biobank), the computing time increases substantially. To complete the analyses within a reasonable timeframe, we used a meta-analysis approach. We performed two sets of meta-analyses, one across two groups within UKBB1 to assess the performance of the meta-analysis, compared to that of the whole UKBB1 data analysis, and the other across UKBB1 and UKBB2.

#### Meta-analyses within UKBB1

We randomly divided the UKBB1 into two groups of equal size (denoted as g1 and g2), and fitted all models mentioned above for each group. P-values from each group were meta-analysed using the Fisher’s method^24^. We then compared these p-values with those based on the whole UKBB1 data set.

#### Meta-analyses across UKBB1 and UKBB2

In UKBB2, 197,394 individuals with genotype data passed the QC, of which 94K have no missing covariates and main response. Similar to meta-analyses within UKBB1, we randomly divided the UKBB2 into two groups of equal size (denoted as G1 and G2), and fitted all models mentioned above for each group. We then meta-analysed the results from G1, G2, and UKBB1 (denoted as G0) using the Fisher’s method^24^. For UKBB2, the same pre-adjustment as for UKBB1 was applied to the main response and covariates as the second trait in MRNM.

#### Software

MTG2: https://sites.google.com/site/honglee0707/mtg2

## ACKNOWLEDGEMENTS

This research is supported by the Australian National Health and Medical Research Council (1080157, 1087889), and the Australian Research Council (DP160102126, FT160100229). This research has been conducted using the UK Biobank Resource. UK Biobank (http://www.ukbiobank.ac.uk) Research Ethics Committee (REC) approval number is 11/NW/0382. Our reference number approved by UK Biobank is 14575.

## AUTHOR CONTRIBUTIONS

S.H.L. conceived the idea and directed the study. G.N. and S.H.L. performed the analyses. N.R.W., J.v.d.W. and E.H. provided critical feedback and key elements in interpreting the results. S.H.L., G.N., N.R.W., J.v.d.W., E.H. and X.Z. drafted the manuscript. All authors contributed to editing and approval of the final manuscript.

